# Network controllability mediates the relationship between rigid structure and flexible dynamics

**DOI:** 10.1101/2021.04.23.441156

**Authors:** Shi Gu, Panagiotis Fotiadis, Linden Parkes, Cedric H. Xia, Ruben C. Gur, Raquel E. Gur, David R. Roalf, Theodore D. Satterthwaite, Danielle S. Bassett

## Abstract

Precisely how the anatomical structure of the brain supports a wide range of complex functions remains a question of marked importance in both basic and clinical neuroscience. Progress has been hampered by the lack of theoretical frameworks explaining how a structural network of relatively rigid inter-areal connections can produce a diverse repertoire of functional neural dynamics. Here, we address this gap by positing that the brain’s structural network architecture determines the set of accessible functional connectivity patterns according to predictions of network control theory. In a large developmental cohort of 823 youths aged 8 to 23 years, we found that the flexibility of a brain region’s functional connectivity was positively correlated with the proportion of its structural links extending to different cognitive systems. Notably, this relationship was mediated by nodes’ boundary controllability, suggesting that a region’s strategic location on the boundaries of modules may underpin the capacity to integrate information across different cognitive processes. Broadly, our study provides a mechanistic framework that illustrates how temporal flexibility observed in functional networks may be mediated by the controllability of the underlying structural connectivity.

**AUTHOR SUMMARY:** Precisely how the relatively rigid white matter wiring of the human brain gives rise to a diverse repertoire of functional neural dynamics is not well understood. In this work, we combined tools from network science and control theory to address this question. Capitalizing on a large developmental cohort, we demonstrated that the ability of a brain region to flexibly change its functional module allegiance over time (i.e., its modular flexibility), was positively correlated with its proportion of anatomical edges projecting to multiple cognitive networks (i.e., its structural participation coefficient). Moreover, this relationship was strongly mediated by the region’s boundary controllability, a metric capturing its capacity to integrate information across multiple cognitive domains.

## INTRODUCTION

The human brain is a complex interconnected system. Neural signals from one region spread to other regions in the system by traveling through underlying nerve fibers. Conceptually, every function has its foundation in structure (Huntenberg, Bazin, & Margulies, 2018; Park & Friston, 2013). Yet in practice, it is methodologically challenging to construct interpretable and theoretically justified relations between structure and function (Suarez, Markello, Betzel, & Misic, 2020). A key challenge lies in addressing precisely how individual neural circuits interact with each other and thereby give rise to brain dynamics. A second challenge lies in addressing the marked differences in structural and functional connectivity fingerprints across individuals (Finn et al., 2015; S. Mueller et al., 2013).

A promising approach to meet these challenges is network neuroscience. Network neuroscience utilizes graph theory to understand connectivity patterns in neural systems. The computational toolbox of network neuroscience can be used to encode data acquired from multiple imaging modalities including diffusion weighted imaging (DWI) and functional magnetic resonance imaging (fMRI) (Bassett & Sporns, 2017). The former allows us to quantify the **structural connectivity** defined from axonal projections, whereas the latter provides time series of the brain’s blood-oxygen-level-dependent (BOLD) activity that can be used to assess its **functional connectivity** (Bassett et al., 2011; Bullmore & Sporns, 2009; Hutchison et al., 2013). Therefore, network neuroscience equips us with a highly appropriate framework to address our question on how the diverse functional expression of the human brain emerges from its underlying structural architecture.

Although several prior studies have utilized graph theoretical principles to predict functional connectivity given the underlying structural connectivity, it still remains unclear how relatively rigid anatomical networks give rise to functional time series encoding flexible dynamic signals (Deco, Jirsa, & McIntosh, 2011; Goñi et al., 2014; Hermundstad et al., 2013; Honey et al., 2009; Mišić et al., 2016; Park & Friston, 2013; Supekar et al., 2010; Vasquez-Rodriguez et al., 2019). An early attempt to tackle this problem focused on the statistical similarity between spatio-temporal structural and functional connectivity patterns (Honey et al., 2009), and reported that functional patterns, although variable, are constrained by the underlying structure. More recent studies have used principles from communication theory to predict activity (Goñi et al., 2014), as well as to suggest that the transient nature of functional connectivity depends on both the anatomical connections and the dynamic coordination of polysynaptic pathways (Shen, Hutchison, Bezgin, Everling, & McIntosh, 2015). Complementary studies have also begun to evaluate the relation between flexible functional expression and enhanced cognitive performance (Baum et al., 2017, 2020; Bertolero, Yeo, & D’Esposito, 2015; Cocuzza, Ito, Schultz, Bassett, & Cole, 2020; Cole, Ito, Cocuzza, & Sanchez-Romero, 2021; Hermundstad et al., 2013; Park & Friston, 2013; Rosenberg, Martinez, et al., 2020; Rosenberg, Scheinost, et al., 2020; Sanchez-Alonso, Rosenberg, & Aslin, 2021; Supekar et al., 2010; Wendelken et al., 2017; Yoo et al., 2020).

A crucial consideration when attempting to bridge structure and function is the identification of descriptive statistics of network organization that can be translated from structural to functional modalities (Cabral, Kringelbach, & Deco, 2017; Murphy, Bertolero, Papadopoulos, Lydon-Staley, & Bassett, 2020). One such descriptive statistic considered in this work is **modular flexibility** (Khambhati, Sizemore, Betzel, & Bassett, 2018). Modular flexibility represents how frequently brain regions change the functional modules they belong to, across time. On a dynamic functional network, a region that is more likely to be connected to multiple functional modules at different time-points is hence more flexible. Indeed, modularity has been reported to flexibly vary across time in a manner that tracks cognitive processes, including working-memory performance (Pedersen, Zalesky, Omidvarnia, & Jackson, 2018), executive function (Baum et al., 2017), learning capability (Bassett et al., 2011), attention (Shine, Koyejo, & Poldrack, 2016), ability to respond to environmental uncertainty (Kao et al., 2020), and overall cognitive flexibility (Braun et al., 2015).

How does this flexible modularity arise from the relatively fixed white matter connectivity patterns? To answer this question, we first turn to the way in which brain regions connect across modules. We use a measure called the **participation coefficient**, which quantifies the relative distribution of a node’s edges between its own and different modules across the full brain network (Baum et al., 2017; Guimera & Amaral, 2005; Power, Schlaggar, Lessov-Schlaggar, & Petersen, 2013). A lower participation coefficient indicates that a node has edges primarily restricted to its own structural or functional **community**, whereas a larger participation coefficient indicates that a node has edges uniformly distributed across multiple communities (Power et al., 2013). To deepen our understanding, we next turn to a measure called boundary controllability, recently introduced in the network control theory (NCT) literature (Pasqualetti, Zampieri, & Bullo, 2014). The structural network architecture of a system, particularly as measured by its **controllability**, can determine the range of dynamics that the system can support (Towlson et al., 2018; Yan et al., 2017). Intuitively, structural boundary controllability measures the necessary input required by a node to drive the overall system along a desired trajectory (Gu, Pasqualetti, et al., 2015). The role of controllability in regulating dynamic brain state transitions as well as predicting the maturity of adolescent brain systems during development has been corroborated by several recent studies (Cornblath et al., 2020; Cui et al., 2020; Tang et al., 2017).

In the present work, we sought to understand the relationship between rigid structure and flexible dynamics. Our approach was to examine the relationships between the participation coefficients of both structural and functional networks, the boundary controllability of structural networks, and the modular flexibility of dynamic functional networks, across regions and among individuals. We hypothesized that the participation coefficients of the structural network architecture would positively correlate with the corresponding flexibility of the functional network. Such a potential association is conceptually justified: a node with edges connecting only to other nodes within its own module would not typically be expected to suddenly alter its connectivity patterns and establish connections with nodes from different modules, over a short period of time. Moreover, we theorized that this relationship between structural participation coefficients and modular flexibility would be fully mediated by the serial effects of boundary controllability and functional participation coefficients. By bridging structure to dynamics in this step-wise fashion, we conceptually unpack the transfer function from rigidity to flexibility.

## MATERIALS AND METHODS

### Data acquisition and pre-processing

We used T1-weighted, diffusion tensor imaging (DTI) and resting-state fMRI BOLD data taken from 823 healthy individuals from the Philadelphia Neurodevelopmental Cohort (PNC) (Ingalhalikar et al., 2014; Satterthwaite et al., 2014). All participants were between 8-23 years of age and their accompanying DTI and fMRI data passed stringent quality control (Roalf et al., 2016; Rosen et al., 2018; Satterthwaite et al., 2014, 2013). All MRI scans were acquired on the same 3T Siemens Tim Trio whole-body scanner with a 32-channel head coil at the Hospital of the University of Pennsylvania. The Institutional Review Boards of both the University of Pennsylvania and the Children’s Hospital of Philadelphia have approved the study procedures.

The standardized structural imaging protocol included a T1-weighted scan obtained using a magnetization-prepared, rapid-acquisition gradient-echo sequence (repetition time: TR = 1810ms, echo time: TE = 3.5ms, field of view: FoV = 180 × 240mm^2^, voxel dimensions = 1 x 1 x 1mm^3^, flip angle = 9°) and a DTI scan acquired using a twice-refocused spin-echo single-shot echo-planar imaging sequence (TR = 8100ms, TE = 82ms, FoV = 240 x 240mm^2^, voxel dimensions = 2 x 2 x 2mm^3^, flip angle = 90°*/*180°*/*180°). The T1-weighted scans were pre-processed using the automated *FreeSurfer* software suite (version 5.3) (Dale, Fischl, & Sereno, 1999; Fischl, Sereno, & Dale, 1999) and parcellated into 234 individualized network nodes based on the Lausanne atlas (Cammoun et al., 2012). Each node was then assigned to one of eight pre-defined functional modules (Yeo et al., 2011): visual, somatomotor, dorsal attention, ventral attention, limbic, frontoparietal control, default mode network, and subcortical. The DTI scans were pre-processed using *FSL*, including skull stripping as well as correction for eddy currents and in-scanner motion (Smith et al., 2004). Deterministic tractography was then implemented using *DSI Studio*, and symmetric adjacency matrices were generated for each subject where the edge weight between two given nodes was defined as the mean **fractional anisotropy** along the connecting streamlines (Yeh, Verstynen, Wang, Fernández-Miranda, & Tseng, 2014). A more detailed description of the parameters used in the proposed processing pipeline can be found in Ref. (Baum et al., 2017).

Resting-state fMRI scans were also acquired for each subject using a BOLD sequence (TR = 3000ms, TE = 32ms, FoV = 192 x 192mm^2^, voxel dimensions = 3 x 3 x 3mm^3^, flip angle = 90°). The functional images were pre-processed using a previously validated pipeline (Ciric et al., 2017). Steps included correction for distortions induced by magnetic field inhomogeneities, removal of the initial four volumes of each acquisition to allow for steady-state magnetization, re-alignment of all volumes to a reference volume, co-registration of functional data to structural data, temporal band-pass filtering, and de-noising (confound regression applied, including 36 regressors as well as spike regression) of the BOLD time series (Ciric et al., 2017; Satterthwaite et al., 2013). In-scanner head motion was defined as the mean relative root-mean-squared displacement calculated during the time series re-alignment step of the pipeline (Satterthwaite et al., 2013). After the data were pre-processed, the scans were parcellated into the same 234 individualized network nodes as the DTI scans. Functional connectivity matrices were finally generated for each subject where the edge weight between two given nodes was defined as the Pearson’s correlation coefficient between their corresponding BOLD signals. The overall pipeline is schematically illustrated in Figure 1.

**Figure 1.**
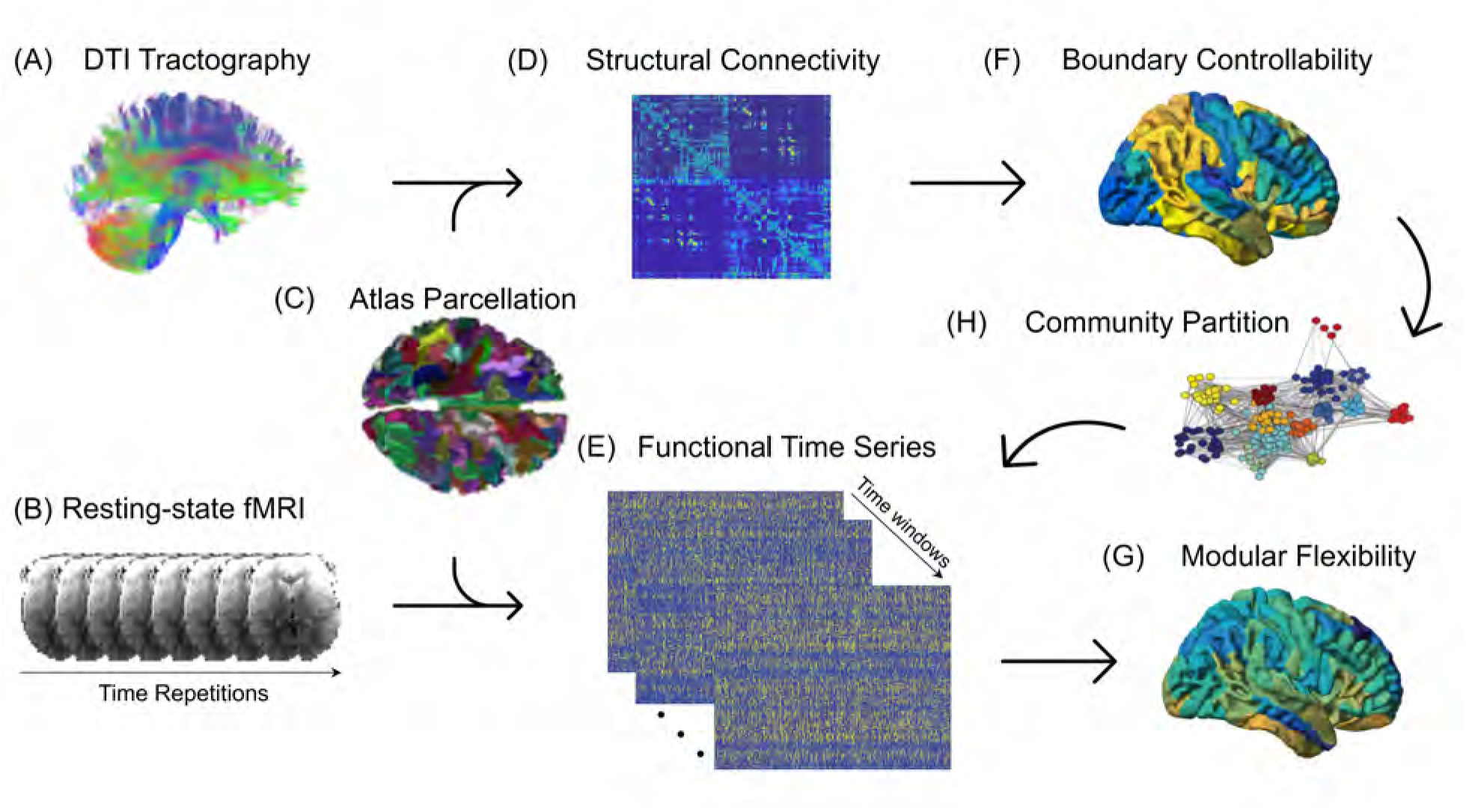
Structural and Functional Processing Pipeline Schematic. The *(A)* diffusive fiber tractography obtained from the DTI scans and *(B)* the resting-state BOLD fMRI time series are parcellated using *(C)* the 234-region Lausanne atlas, to construct *(D)* structural connectivity matrices and *(E)* functional time series for each subject. The structural connectivity matrices are then used to compute each region’s *(F)* boundary controllability, whereas the functional time series are used to assess the modular dynamics of the time-resolved functional networks by calculating *(G)* the modular flexibility of each region. In our proposed analysis, the participation coefficients obtained from *(H)* the community partition of the static networks act as mediators in predicting how flexible the functional network will be, given its structural connections, thereby bridging the two imaging modalities.

### Participation Coefficients

Participation coefficients measure the extent to which a node’s connectivity profile participates diversely across modules. Mathematically, given a network wherein *N*_*m*_ designates the total number of modules considered (here set to eight), *s* iterates through the eight pre-defined functional modules mentioned in the previous section, 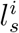 represents the number of links between node *i* and nodes in module *s*, and 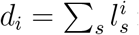 represents the total degree of node *i*, the participation coefficient of node *i* is defined as:

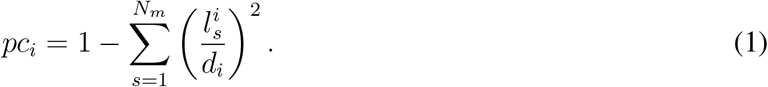

Participation coefficients range between zero and one, where a value of zero indicates that a node’s edges are entirely restricted to its own community and a value of one indicates that a node’s edges uniformly extend across all other modules in the network (Power et al., 2013).

### Boundary Controllability of Structural Networks

NCT is a mathematical framework that aims to assess whether a network can be controlled. Specifically, NCT asks whether the output of the overall system can be driven towards a desired outcome given a set of input signals. There are several metrics from NCT that attempt to quantify a node’s overall ability to alter other nodes’ neurophysiological states (Pasqualetti et al., 2014). Here, we focused on a metric called boundary controllability. Assuming we have a structural connectivity matrix constructed from a DTI scan, the boundary controllability of a node is a heuristic metric predicting its ability to integrate information across different cognitive processes. In other words, brain regions with high boundary controllability tend to lie at the boundaries between network communities, and are thus thought to be structurally predisposed to efficiently control the integration of different cognitive systems (Gu, Pasqualetti, et al., 2015).

In order to calculate the boundary controllability of each brain region, we first partitioned the cortical mantle into communities, using a common community-detection algorithm (Louvain-like locally greedy heuristic algorithm) (Bassett et al., 2013; Blondel, Guillaume, Lambiotte, & Lefebvre, 2008; Gu, Pasqualetti, et al., 2015). Based on that community partition, we identified an initial set of boundary nodes (*N*_1_) and assigned them a boundary controllability value of one (Pasqualetti et al., 2014). Then, we used an iterative process to further partition the network into communities to identify more boundary nodes at increasingly finer levels of the modular hierarchy, until all nodes were assigned a boundary controllability value. During each step of this iterative process, the boundary controllability value of each node *i* was set to (*N* − *N*_*i*_)*/N*, where *N* is the total number of cortical brain regions and *N*_*i*_ is the number of nodes on the boundary between communities (Gu, Pasqualetti, et al., 2015).

### Modular Flexibility of Functional Networks

Brain regions have been shown to interact among themselves across multiple temporal scales, even at rest (Betzel et al., 2019; Meunier, Lambiotte, Fornito, Ersche, & Bullmore, 2009; M. E. Newman, 2006; Rubinov & Sporns, 2010). This property gives rise to modular dynamics which can be assessed quantitatively by modular flexibility (Bassett et al., 2011).

The parcellated time series of each subject **x**_*N*×*T*_ (N = 234 regions, T = 120 TRs) were divided into 10 non-overlapping time-windows, and the temporal network 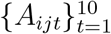 was constructed by defining *A*_*ijt*_ as the Pearson’s correlation coefficient between the BOLD time series of regions *i* and *j* within the *t*^*th*^ sliding window Telesford et al. (2016). Each time-window corresponded to a layer in the multi-layer network, and the multi-layer signed modularity function was defined as (Mucha, Richardson, Macon, Porter, & Onnela, 2010):

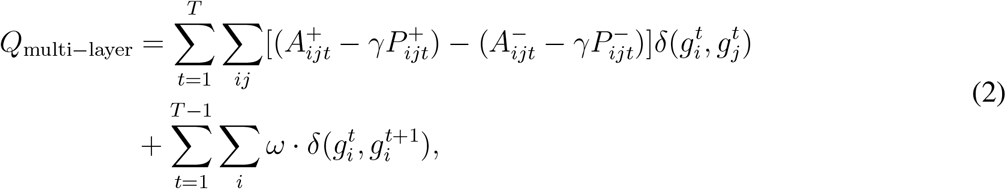

where 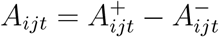 is decomposed into its positive and negative parts with 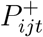 and 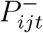 representing the corresponding parts obtained from null models (Newman-Girvan) (M. E. J. Newman & Girvan, 2004). The label 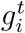 denotes the community assignment of node *i* in the *t*^*th*^ layer, *δ*(*x, y*) is the Kronecker-*δ* function set equal to 1 if *x* = *y* and to 0 otherwise, *γ* is a parameter that tunes the size of communities (here, equal to 1), and *ω* represents the coupling strength between neighboring layers (here, equal to 1). A recent study identified that the test-retest reliability in calculating dynamic network measures, such as modular flexibility, depended on parameter selection (i.e., *γ* and *ω*), among other factors (Yang et al., 2020). Even though that study identified the parameter value pair with the overall highest intra-correlation coefficient to be (*γ, ω*) = (1.05, 2.05), we implemented the more widely used pair (*γ, ω*) = (1, 1) (Bassett et al., 2011; Betzel, Satterthwaitte, Gold, & Bassett, 2017; Braun et al., 2015; Pedersen et al., 2018; Telesford et al., 2016) because the corresponding modular flexibility values were virtually identical (*r* = 0.963, *p* = 0).

In order to explore the temporal evolution of each module in the multi-layer network, each node within each time-window was assigned into a community, indicating its module allegiance. For this purpose, a Louvain-like locally greedy heuristic algorithm (Braun et al., 2015) was used to maximize the modularity index *Q*_multi−layer_. This process gave rise to a partition matrix **G**_*N*×*T*_ (N = 234 regions, T = 10 time-windows) in which *G*_*i,t*_ represented the community to which node *i* in layer *t* belonged. The nodal flexibility *f*_*i*_ of each region *i* was then defined as:

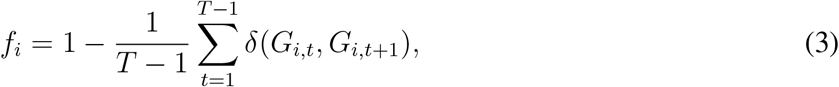

and assesses how often brain region *i* shifts its community assignment between temporal layers. On the subject level, the global flexibility of the entire network was defined as the mean of all regional *f*_*i*_ values: 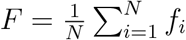. An in-depth description of the method can be found in Ref. (Khambhati et al., 2018).

### Statistical Analyses

All analyses were performed using the SPSS statistical software (v26, Armonk, NY: IBM Corporation) and Python (v3.9). A threshold for significance of *p <* 0.05 was used. Pearson’s correlation coefficients (*r*) and *p*-values were reported for each bivariate analysis. In order to address the potential spatial autocorrelation between our structural and functional metrics of interest, we applied a previously validated spatial permutation framework (i.e., spin test with 100,000 permutations) to generate null models (Alexander-Bloch et al., 2018). For those analyses (i.e., Figures 2A and D), the corresponding *p*-values are reported as *p*_*spin*_. Moreover, for all multiple linear regression analyses performed, age, sex, and in-scanner head motion were adjusted for, and correction for multiple comparisons was performed using the Benjamini-Hochberg false discovery rate (FDR) procedure (i.e., Figures 4 and 5) (Benjamini & Hochberg, 1995).

**Figure 2.**
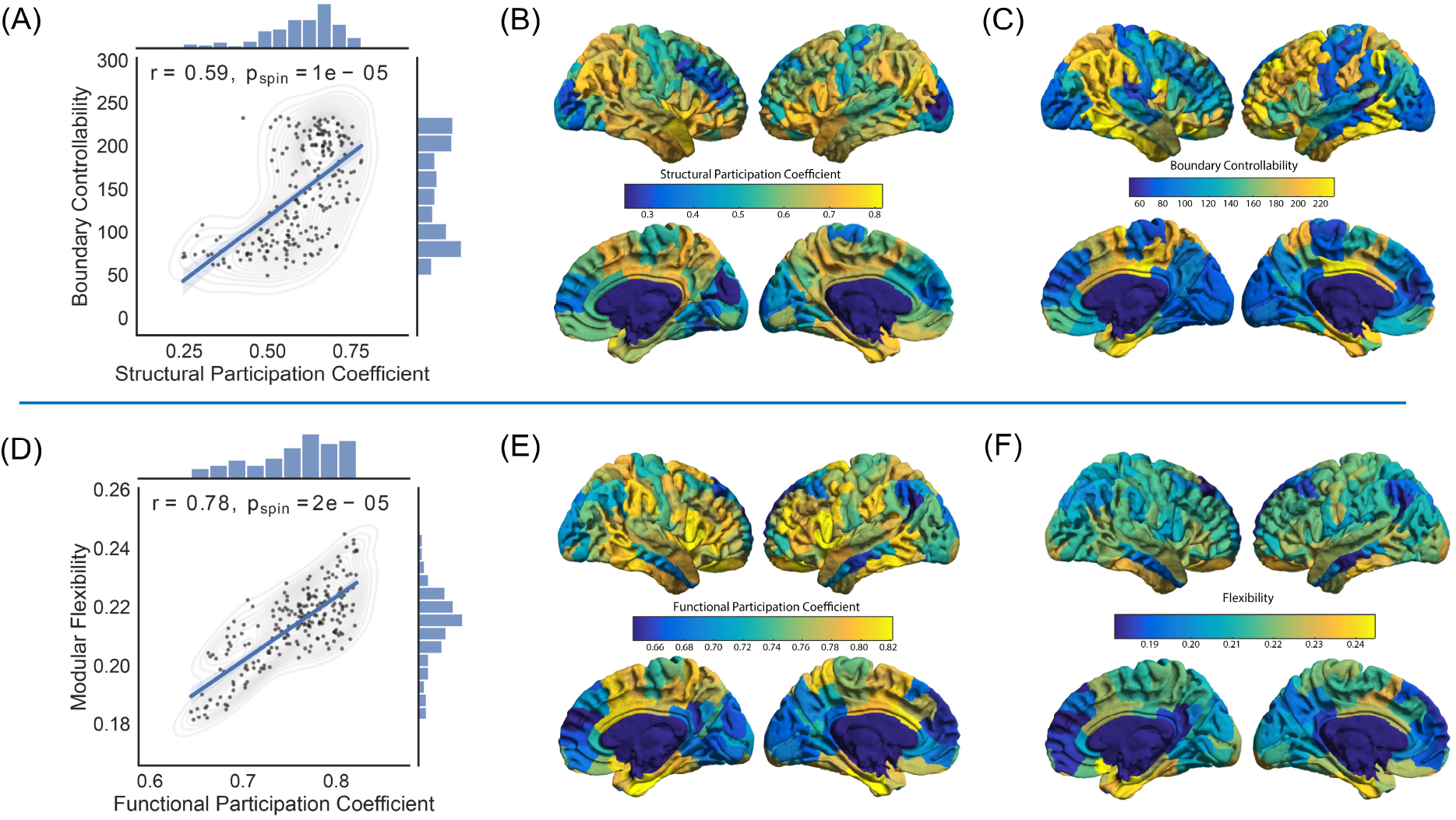
Region-wide patterns. Structural patterns are shown in the first row: *(A)* Boundary controllability was significantly correlated with the structural participation coefficient (*r* = 0.59, *p*_*spin*_ = 1 × 10^−5^; the shaded areas correspond to the confidence curves for the fitted line). In order to address the potential spatial autocorrelation between boundary controllability and the structural participation coefficient, we applied a spatial permutation framework (spin test; see Methods). We also show the average patterns of regional fluctuations of *(B)* the structural participation coefficients and *(C)* boundary controllability, across all subjects. Functional patterns are shown in the second row: *(D)* Modular flexibility was significantly correlated with the functional participation coefficient (*r* = 0.78, *p*_*spin*_ = 2 × 10^−5^). In order to address the potential spatial autocorrelation between modular flexibility and the functional participation coefficient, we applied a spatial permutation framework (spin test; see Methods). We also show the average patterns of regional fluctuations of *(E)* the functional participation coefficients and *(F)* modular flexibility across all subjects.

### Mediation Model

All mediation analyses were performed using the *PROCESS* (v3.4) statistical macro for SPSS (Hayes, 2017). Structural participation coefficients were designated as the independent variable, boundary controllability as the first mediator, functional participation coefficients as the second mediator, and modular flexibility as the dependent variable. Age, sex, and in-scanner head motion were used as covariates in all subject-wide analyses. The hypothesized serial mediation effect was tested using bootstrapping (10,000 samples). Mediation was deemed significant if the bootstrapping confidence interval did not include zero. Unstandardized regression coefficients (*c*) and *p*-values were reported for each association within the mediation analysis.

For the purpose of maintaining consistency, all variables within the mediation model were rescaled to range from zero to one. Moreover, in order to incorporate temporal directionality into the model, the functional participation coefficient of each node was calculated only during the first time-window of the resting-state fMRI BOLD sequence, whereas its modular flexibility was averaged across all remaining time-windows (second through tenth).

## RESULTS

### Structural Participation Coefficients Positively Correlate with Boundary Controllability

A brain region’s structural participation coefficient quantifies its role in communicating across multiple modules (Hall et al., 2018). Similarly, boundary controllability assesses a region’s predicted ability to integrate information from different cognitive modules, attributing higher values to regions that are located on the boundary of larger modules (Medaglia et al., 2018). Therefore, we hypothesized that these two metrics would be positively correlated. To examine this hypothesis, we computed both metrics for each region and averaged them across all subjects. We observed a strong positive correlation (*r* = 0.59, *p*_*spin*_ = 1 × 10^−5^) between the structural participation coefficient and boundary controllability across regions (Figure 2A). Regionally, the parietal and temporal lobes displayed high values of the participation coefficient and boundary controllability, whereas the occipital lobe displayed low values (Figures 2B and C).

### Functional Participation Coefficients Positively Correlate with Modular Flexibility

As described earlier, modular flexibility assesses how often a node shifts its community assignment across different time-windows. Functional participation coefficients reflect the same property on the static level; that is, during one time-window. Indeed, some regions known as connector hubs, play a gating role across multiple communities and have a larger functional participation coefficient (Bertolero, Yeo, Bassett, & D’Esposito, 2018; Cohen & D’Esposito, 2016). The same regions would also be theoretically expected to often shift their allegiance between different cognitive networks, across multiple temporal scales. Thus, we hypothesized that the participation coefficients obtained from the static functional network would be positively correlated with the functional network’s flexibility across time. To test our hypothesis, we first computed these two metrics for each region and averaged the values across subjects. We observed that the functional participation coefficients and the modular flexibility were strongly positively correlated across regions (*r* = 0.78, *p*_*spin*_ = 2 × 10^−5^; Figure 2D). Moreover, the temporal lobe displayed high values of functional participation coefficient and modular flexibility, whereas the medial frontal lobe displayed low values (Figures 2E and F).

### Boundary Controllability and Functional Participation Coefficients Serially Mediate the Relationship between Structural Participation Coefficients and Modular Flexibility

In the previous two sections, we established that a region’s ability to dynamically interact with multiple cognitive modules (via structural controllability and functional flexibility) was also reflected on the static level (via participation coefficients), in both structural and functional modalities. We next attempted to bridge the two imaging modalities. We hypothesized that the structural participation coefficient of a region would predict its functional modular flexibility via the serial mediation effects of its boundary controllability and functional participation coefficient. In testing this hypothesis, we found that, across regions, the structural participation coefficient (the independent variable) was positively correlated with temporal modular flexibility (the dependent variable) (*r* = 0.16, *p*_*spin*_ = 0.012). As theorized, this effect was serially mediated by boundary controllability and functional participation coefficients (Figure 3; total effect = 0.215; *p* = 0.002, indirect effect = 0.190; Bootstrapping Confidence Interval = [0.119 0.269]).

**Figure 3.**
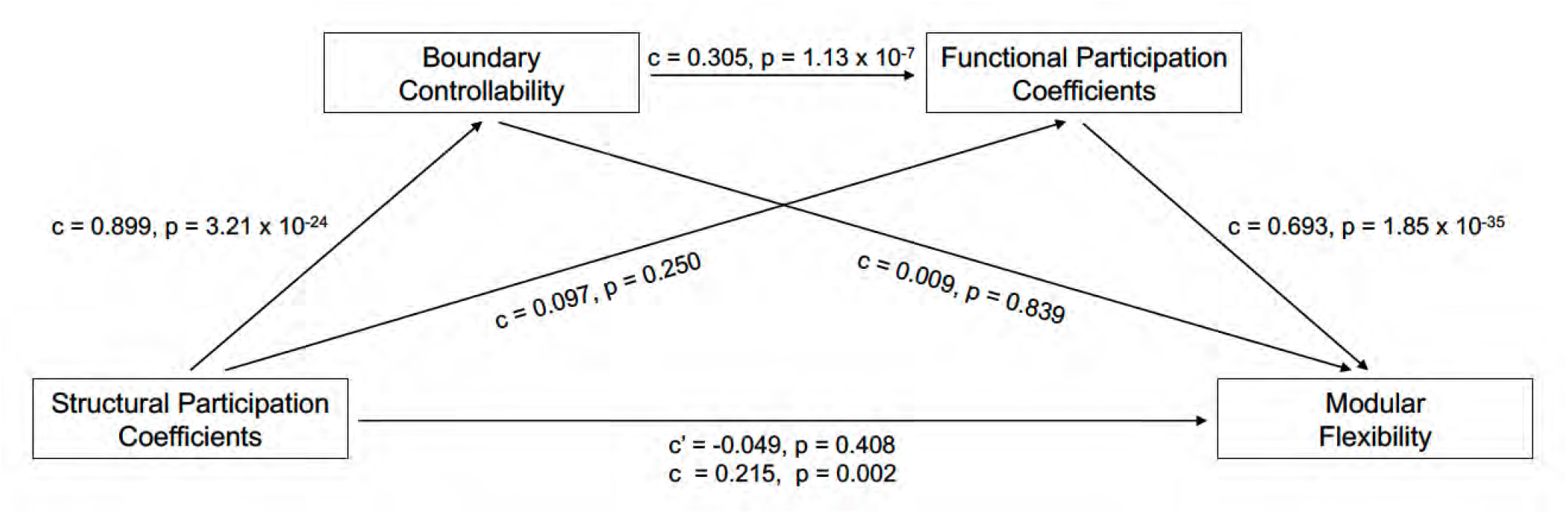
Serial Mediation Model. We examined the hypothesis that the structural participation coefficient of each brain region (obtained from the DTI sequences) can predict its modular flexibility (obtained from the resting-state fMRI BOLD sequences) via the serial mediation effects of boundary controllability and the functional participation coefficient. In order to establish temporal directionality within the mediation model, the functional participation coefficient was calculated only during the first time-window of the functional time series and modular flexibility was averaged across all remaining time-windows (second through tenth). The unstandardized regression coefficient (*c*) and *p*-value are reported for each association within the mediation analysis. Moreover, the total and direct effects of the structural participation coefficient (independent variable) on modular flexibility (dependent variable) are also provided (total effect: regression coefficient *c, p*-value; direct effect: regression coefficient *c*′, *p*-value).

**Figure 4.**
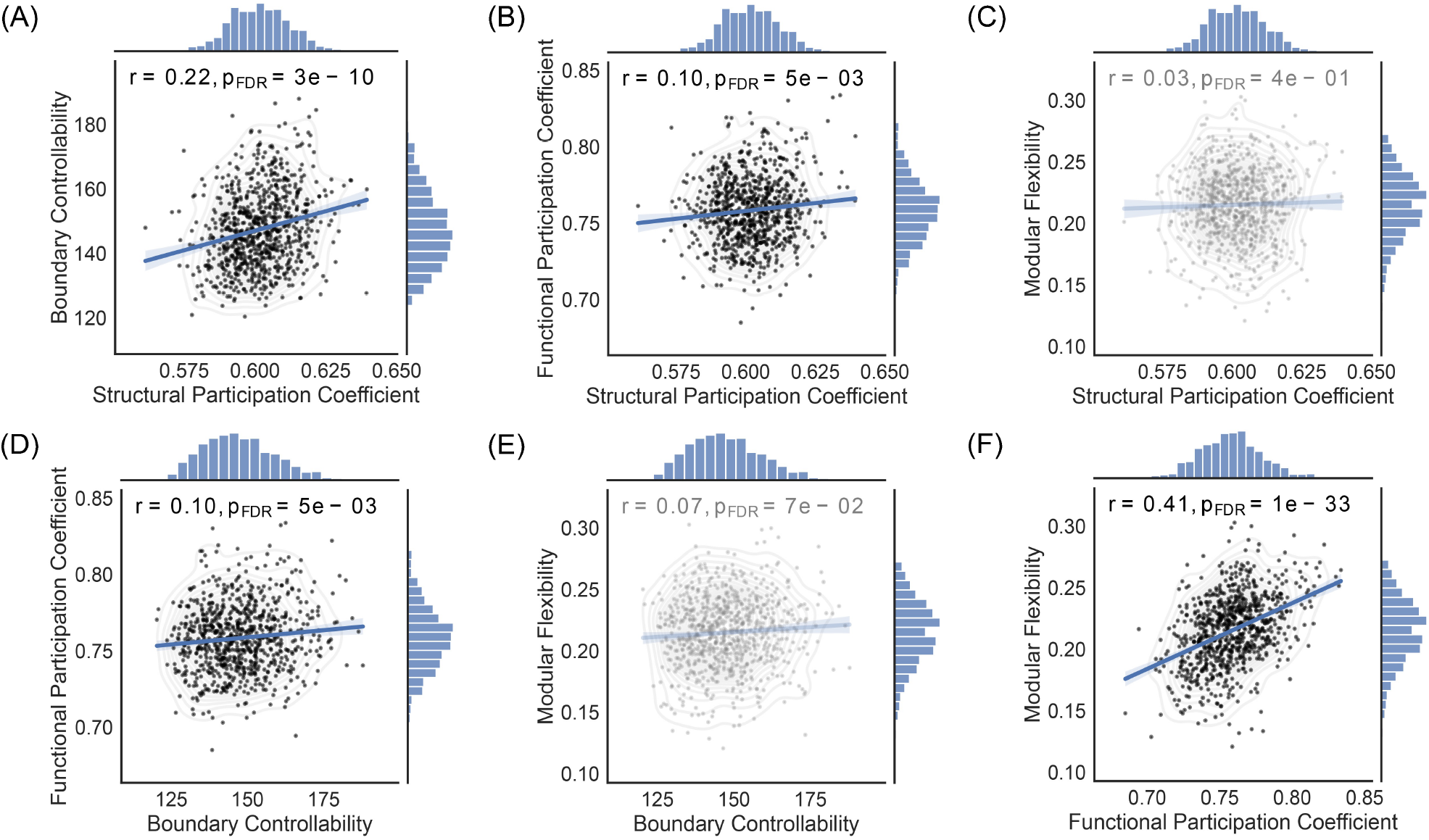
Subject-wide correlations. We examined whether the correlations observed within our structural and functional variables across regions persisted across subjects. In all analyses, we adjusted for age, sex, and in-scanner head motion (FDR corrected for multiple comparisons). Here we show leverage plots corresponding to the multiple linear regression models used: *(A)* The structural participation coefficient was significantly correlated with boundary controllability (*r* = 0.22, *p* = 3 × 10^−10^) and with *(B)* the functional participation coefficient (*r* = 0.10, *p* = 0.005), but not with *(C)* modular flexibility (*r* = 0.03, *p* = 0.40). *(D)* Global boundary controllability was also positively associated with the average functional participation coefficient for each subject (*r* = 0.10, *p* = 0.005) and had a trending relationship with *(E)* modular flexibility (*r* = 0.07, *p* = 0.07). Lastly, as in the regional case, *(F)* each subject’s average functional participation coefficient was strongly correlated with its corresponding modular flexibility (*r* = 0.41, *p* = 1 × 10^−33^). The shaded areas correspond to the confidence curves for the fitted lines.

**Figure 5.**
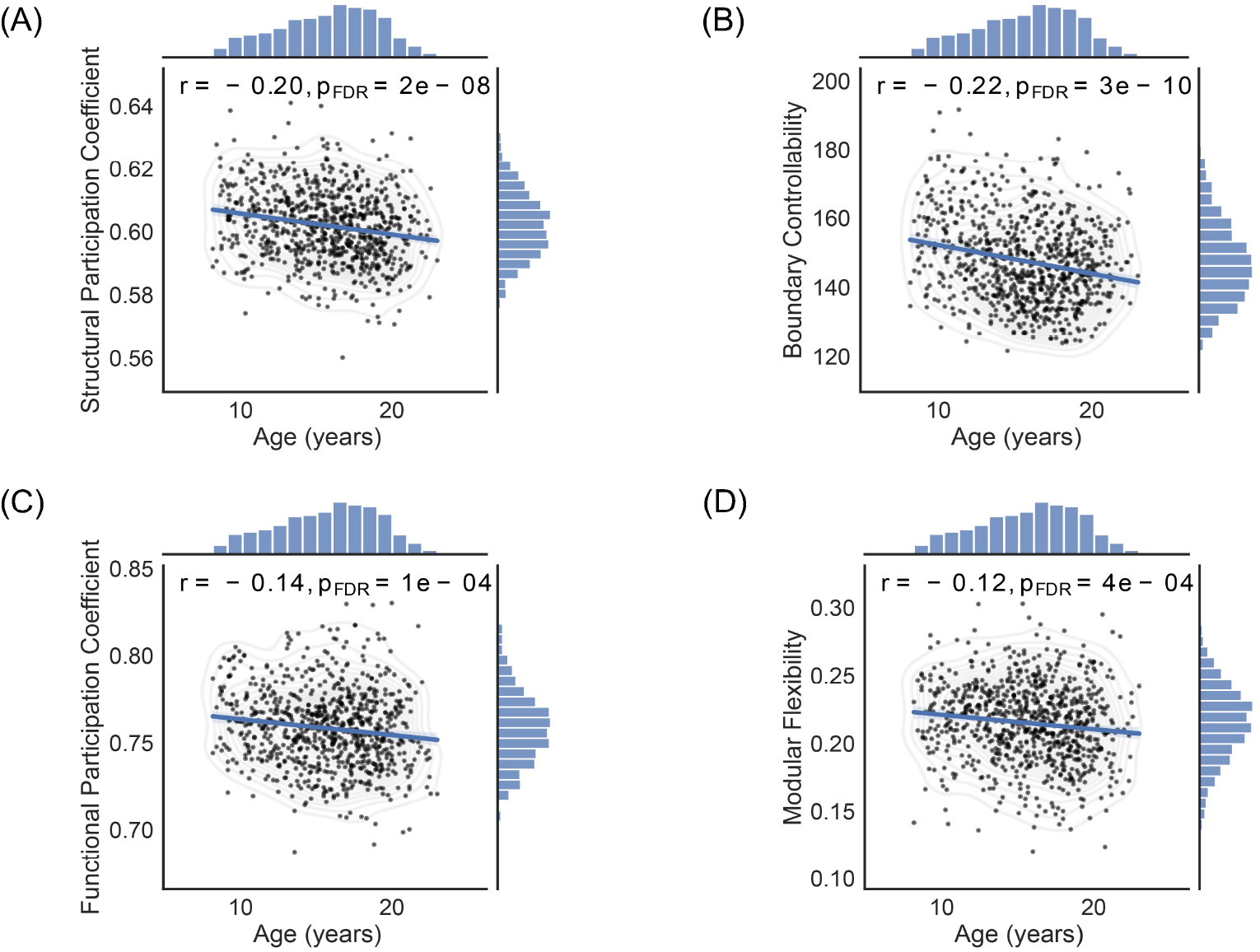
Age-related changes in structural and functional metrics. Global structural and functional metrics were derived per individual by averaging the corresponding values across brain regions. *(A)* The structural participation coefficient (*r* = −0.20, *p* = 2 × 10^−8^; FDR), *(B)* boundary controllability (*r* = −0.22, *p* = 3 × 10^−10^; FDR), *(C)* functional participation coefficient (*r* = −0.14, *p* = 1 × 10^−4^; FDR), and *(D)* modular flexibility (*r* = −0.12, *p* = 4 × 10^−4^; FDR) all declined linearly with age. The effects of sex and in-scanner head motion were regressed out, and each analysis was FDR corrected for multiple comparisons. The shaded areas correspond to the confidence curves for the fitted lines.

It is noteworthy that the significant positive correlation between structural participation coefficients and (the second mediator) functional participation coefficients (*r* = 0.32, *p* = 3.88 × 10^−7^) became non-significant after regressing out (the first mediator) boundary controllability (*c* = 0.097; *p* = 0.25). Similarly, even though the positive association between boundary controllability and modular flexibility was significant (*r* = 0.32, *p* = 5.54 × 10^−7^), it became non-significant once the functional participation coefficient was regressed out of the model (*c* = 0.009, *p* = 0.84). More importantly, however, the direct effect of the structural participation coefficient on modular flexibility became non-significant once the effects of the mediators were regressed out of the model (Figure 3; *c* = −0.049, *p* = 0.41). These statistical results strongly support the notion that the total effect of the structural participation coefficient on modular flexibility is indeed fully mediated by the serial effects of boundary controllability and the functional participation coefficient.

### Relationship between Participation Coefficients, Boundary Controllability, and Modular Flexibility, across subjects

Thus far, we have focused on regional variation in structure and dynamics, and their relations. Next, we turn to subject-level variation to better understand inter-individual differences. Specifically, we examine the relationship(s) between participation coefficients, boundary controllability, and modular flexibility, across subjects. We used multiple linear regression models to adjust for age, sex, and in-scanner head motion (corrected for multiple comparisons using FDR). As in the regional analyses, we observed that the structural participation coefficient was positively correlated with boundary controllability (Figure 4A; *r* = 0.22, *p* = 3 × 10^−10^; FDR) and with the functional participation coefficient (Figure 4B; *r* = 0.10, *p* = 0.005; FDR), but not with modular flexibility (Figure 4C; *r* = 0.03, *p* = 0.40; FDR). Moreover, boundary controllability was correlated with the functional participation coefficient (Figure 4D; *r* = 0.10, *p* = 0.005; FDR) and had a trending association with modular flexibility (Figure 4E; *r* = 0.07, *p* = 0.07; FDR). Lastly, we observed a strong positive correlation between average functional participation coefficient and modular flexibility (Figure 4F; *r* = 0.41, *p* = 1 × 10^−33^; FDR). We note that in each aforementioned multiple regression model, age was significantly and consistently correlated with the dependent variable (Figure 4A: *p* = 1.6 × 10^−7^, Figure 4B: *p* = 0.002, Figure 4C: *p* = 0.003, Figure 4D: *p* = 0.002, Figure 4E: *p* = 0.007, and Figure 4F: *p* = 0.066; FDR).

Although we observed no significant correlation between the structural participation coefficient and modular flexibility across subjects, a mediation effect between the two variables could still exist (Hayes, 2018). Thus, we examined whether boundary controllability and the functional participation coefficient still played a serial mediation effect in regulating the relationship between the structural participation coefficient and modular flexibility. Once again, we observed a significant mediation effect in the same direction as observed in the regional analyses (total effect = 0.034, *p* = 0.37; indirect effect = 0.006; Bootstrapping Confidence Interval = [0.0002 0.0125]). However, when age, sex, and in-scanner head motion were also included as covariates in the mediation model, the significance disappeared (total effect = -0.005, *p* = 0.90; indirect effect = 0.003; Bootstrapping Confidence Interval = [-0.001 0.009]).

### Developmental Trajectories of Participation Coefficients, Boundary Controllability, and Modular Flexibility

Due to the substantial effects of age in the aforementioned subject-wide analyses, we tracked the age-related changes in all pertinent variables. We specifically examined the relationships between age and the structural and functional participation coefficients, boundary controllability, and modular flexibility, after regressing out the effects of sex and in-scanner head motion (corrected for multiple comparisons; FDR). Notably, all variables of interest decreased linearly with age, with the structural markers (i.e., structural participation coefficient, Figure 5A, *r* = −0.20 and *p* = 2 × 10^−8^; boundary controllability, Figure 5B, *r* = −0.22 and *p* = 3 × 10^−10^) displaying larger effect sizes than their functional counterparts (i.e., functional participation coefficient, Figure 5C, *r* = −0.14 and *p* = 1 × 10^−4^; modular flexibility, Figure 5D, *r* = −0.12 and *p* = 4 × 10^−4^).

## DISCUSSION

The brain is an interconnected dynamical system whose functional expression relies on the underlying white matter architecture (Deco et al., 2013). The intrinsic mechanisms, however, of how such a diverse repertoire of functions emerges from a relatively rigid anatomical backbone have yet to be fully understood. In this study, we combined tools from network neuroscience and control theory to examine whether and how white matter tractography networks support the flexible modular architecture of the brain, as derived from resting-state fMRI BOLD signals. In structural networks, we found that a region’s participation coefficient was strongly positively correlated with its boundary controllability, suggesting that its ability to be controlled by external input can be assessed by the distribution of edges within its own and different modules across the network. Similarly, in functional networks, we found that a region’s participation coefficient strongly correlated with its modular flexibility, suggesting that a region’s flexibility across multiple temporal windows can be captured by its static participation in the communication occurring both within and between modules. Collectively, these observations provide us with foundational intuitions regarding how temporally-evolving patterns of communication can arise from fixed structural connectomes.

### Structure-function relations across anatomical regions

Once a network structure is provided, one can begin to understand the anatomical support for various patterns of communication both within and between its component modules (Avena-Koenigsberger, Misic, & Sporns, 2018). Each module (or community) consists of a group of densely interconnected nodes (Sporns & Betzel, 2016). Each node can then be assigned a participation coefficient which quantifies its connectivity distribution across communities (Guimera & Amaral, 2005). A high participation coefficient indicates strong between-module and weak within-module connectivity; a low participation coefficient indicates a more uniformly distributed connectivity pattern across modules. Intuitively, whether a region’s connections remain local to or expand beyond their community could partially determine that region’s control over signal transduction throughout the network (Gu, Pasqualetti, et al., 2015; Medaglia et al., 2018). Here we validated that intuition in our structural analyses, where the participation coefficient of a given node displayed a strong correlation with its boundary controllability. The result also deepens our understanding of the brain’s structural organization. The participation coefficient is calculated from a single scale of modules as defined by the eight cognitive systems examined (Yeo et al., 2011), whereas the boundary controllability assesses a region’s location betwixt modules defined across all scales, from eight modules to *N* modules. Hence, the strong relation between these two variables indicates that submodules tend to be formed by larger modules breaking at a hinge, rather than by the center of a large module falling out, like a donut hole. It would be interesting in future work to further examine individual differences in these hinge-like hierarchies and assess their relevance for cognitive function.

Whereas white matter structure provides anatomical support for communication patterns, the brain’s functional dynamics can provide more direct measurements of those putative patterns. In a functional brain network, the participation coefficient can be used to assess how uniformly the edges of a node span modules in a single temporal window, which is typically chosen to be the full duration of the functional scan (Power et al., 2013). In contrast, modular flexibility can be used to assess how the allegiance of a node to a module changes over multiple temporal windows, or over different time scales (Khambhati et al., 2018). For instance, a node that constantly shifts its allegiance between different communities over a task duration or resting-state period would have a high modular flexibility value. Flexibility is a fundamental property of dynamical and adaptive systems, which is thought to support a range of human cognitive processes including motivation (O’Ralley, 2020), working memory (Pedersen et al., 2018), and cognitive flexibility (Braun et al., 2015; Ramos-Nunez et al., 2017), is age-dependent (Malagurski, Liem, Oschwald, Merillat, & Janche, 2020; Schlesinger, Turner, Lopez, Miller, & Carlson, 2017), and can be modulated by mood (Betzel et al., 2017), exercise (Sinha, Berg, Yassa, & Gluck, 2021), and hormonal fluctuations (J. M. Mueller et al., 2021). Following existing literature discussing the relationship between static and dynamic connectivity (Betzel, Fukushima, He, Zuo, & Sporns, 2016), we asked whether a region’s static participation coefficient was associated with its dynamic flexibility. That is, if a region is – on average – strongly connected to multiple modules, does that region also have a propensity to change the module to which it is most strongly connected over short time periods? We find that that answer is “yes”: a region’s functional participation coefficient and modular flexibility were strongly positively correlated. This correspondence between the temporal average and the time-resolved behavior provides deeper insight into the nature of regional roles within a functional network. What appears over all time-windows as a broad participation is in fact produced by temporally resolved affiliations, where a region participates most closely with different modules at different time-points.

### What mediates the relationship between structure and dynamics?

Given the cognitive and clinical relevance of flexibility (Bailey, Aboud, Nguyen, & Cutting, 2018; Barbey, 2018; Bassett, Yang, Wymbs, & Grafton, 2015; Chong et al., 2019; Doucet, Bassett, Yao, Glahn, & Frangou, 2017; Finc et al., 2017; Harlalka, Bapi, Vinod, & Roy, 2019; Rolls, Cheng, & Feng, 2021; Shine, Bissett, et al., 2016; Zhang et al., 2016), it is important to establish a statistically principled pathway whereby modular flexibility could be regulated, and might depend upon underlying structure. Accordingly, we tested the hypothesis that a region’s structural participation coefficient predicts its corresponding functional flexibility, via the serial mediation effects of its boundary controllability and functional participation coefficient. In order to account for temporal directionality in this mediation model, we computed each region’s functional participation coefficient only during the first time-window (out of a total of 10 non-overlapping time-windows) of the resting-state fMRI BOLD sequence, and we calculated its modular flexibility across all remaining time-windows of the same sequence. Consistent with our hypothesis, we found a strong serial mediation effect between structural participation coefficients and functional flexibility, which was fully mediated by the serial effects of boundary controllability and functional participation coefficient (Figure 3). This directional mediation effect was strong – yielding a ratio of indirect to total effect of 0.88 – in predicting the functional flexibility of a region given its structural participation coefficient (Kenny, Kashy, & Bolger, 1998). Notably, when the mediation effects of the two mediators were regressed out of the model, the significant association between structural participation coefficients and modular flexibility disappeared, suggesting that a full mediation effect was present (Hayes, 2018; Rucker, Preacher, Tormala, & Petty, 2011).

Overall, this result shows how a static structural property of a region can mediate a dynamic functional property of the same region. Specifically, the distribution of a region’s edges at a given time-point can mediate how a region shifts the edges’ module allegiance over a period of time. Our observation that boundary controllability and functional participation coefficients strongly mediated this relationship, demonstrates that in addition to having edges reaching different modules across the network, a region’s strategic placement on the boundaries of such modules is also important in assessing its flexibility. This dependence is intuitive, and existing on the boundary bestows the region with the ability to potentially integrate information across different cognitive processes. Indeed, connector hubs – nodes with diverse connections across modules and anatomically located at the boundaries between communities – have been previously reported to coordinate connectivity changes occurring between nodes from different communities, during cognitive tasks (Bertolero et al., 2018; Gratton, Laumann, Gordon, Adeyemo, & Petersen, 2016).

### Structure-function relations across individuals & through development

In order to examine how the relationships between participation coefficients, structural controllability, and functional flexibility varied across participants, we computed a global average metric for each structural and functional marker, per individual, and repeated the above analyses across a large developmental cohort of youth (PNC) (Satterthwaite et al., 2014). Overall, the associations between the variables of interest remained significant in multiple linear regression models adjusting for age, sex, and in-scanner head motion, as in the region-wide analyses. There were, however, two main differences between the across-region and the across-subjects results: (i) the relationship between the structural participation coefficient and modular flexibility became non-significant in the across-subjects analysis, and (ii) age was consistently correlated with the dependent variable in all regression models.

Even though there was no correlation between the structural participation coefficient and modular flexibility in the across-subjects analysis, we re-tested our hypothesis that the relationship between the two variables was still mediated by the structural networks’ boundary controllability and the functional networks’ participation coefficients (Hayes, 2018). Similarly to the across-region analysis, we discovered that a mediation effect of the same directionality was still present and significant. Notably, however, when we included age as a covariate, the mediation effect was no longer significant. This observation could reflect the instrumental role that age plays in shaping structural and functional connectivity during this developmental period.

The important effect of age in the subject-wide mediation model, in addition to our earlier observation that age was strongly correlated with each one of the variables explored in the subject-wide analyses, motivated us to study the age-related changes in all structural and functional variables of interest. Such changes are anticipated, given that the age range of our cohort spans childhood and adolescence, which is a critical period of neurodevelopment and neuroplasticity (Brun et al., 2009; Casey, Tottenham, Liston, & Durston, 2005; Foulkes & Blakemore, 2018; Fuhrmann, Knoll, & Blakemore, 2015; Lenroot et al., 2009; Somerville et al., 2018). We found that both sets of structural (i.e., structural participation coefficient and boundary controllability) and functional (i.e., functional participation coefficient and modular flexibility) metrics decreased linearly with age, with the former displaying larger effect sizes.

The robust age-related decrease of the structural participation coefficient across youth observed in this study has been previously reported (Baum et al., 2017; Dosenbach et al., 2010; Fair et al., 2009; Gu, Satterthwaite, et al., 2015). As a region’s participation coefficient decreases, it develops strong within-module connectivity and weak between-module connectivity. The corresponding increase in modular segregation across development has been shown to enhance network efficiency via the strengthening of hub edges, and to support executive performance (Baum et al., 2017). Here we further observe that boundary controllability decreases with age, while other metrics of controllability (average and modal) have been reported to increase over the same developmental period (Tang et al., 2017). This pattern of findings is particularly notable as boundary controllability is driven by changes in modular architecture, whereas neurodevelopmental changes in average and modal controllability are not (Gu, Pasqualetti, et al., 2015; Tang et al., 2017). These observations suggest that a fundamental change in graph architecture is taking place throughout this developmental period that potentially contributes to the maturity of brain modules, in support of the emergence of functional roles of cognitive systems (Gu, Satterthwaite, et al., 2015) associated with network segregation (Baker et al., 2015; Baum et al., 2017; Fair et al., 2009, 2007). Furthermore, the changes in modular flexibility that we observed across this period could represent enhanced communication plasticity, paving the way for the emergence of high-order cognitive functions characteristic of adulthood (Geerligs, Saliasi, Maurits, & Lorist, 2015; Luna, Marek, Larsen, Tervo-Clemmens, & Chahal, 2015).

## LIMITATIONS & FUTURE DIRECTIONS

The work should be examined in light of several methodological considerations and limitations. This study, by design, was cross-sectional aiming to explore how functional flexibility emerged from the underlying white matter architecture. A cross-sectional design is limited in its ability to tease apart temporal precedence. As such, it would be highly informative to also include longitudinal data that address how the structural and functional properties of each region change during this crucial period of neurodevelopment. In order to account for temporal directionality in our mediation analyses, we computed functional participation coefficients from the first temporal window of the fMRI BOLD sequence and modular flexibility from the remaining time-windows. This approach could, however, raise the issue that incorporating functional signals from a single time-window with limited time series length could inflate signal noise (Noble, Scheinost, & Constable, 2019). In order to address this potential concern, we repeated our mediation analyses after computing mean functional participation coefficients and modular flexibility from all time-windows; all conclusions remained the same. Moreover, although most of the structural and functional markers examined here have been separately reported to regulate cognitive functions such as executive function (Baum et al., 2017; Reineberg & Banich, 2016), processing speed (Varangis, Habeck, Razlighi, & Stern, 2019), and working memory (Stevens, Tappon, Garg, & Fair, 2012), it would be beneficial to incorporate clinical and neurocognitive dimensions into our mediation models. Incorporating behavioral data will address the question of how the serial mediation model as a whole could shape behavior and cognition, and how deficits in the inter-relations among its components could potentially lead to neurological and psychiatric developmental disorders (Aerts, Fias, Caeyenberghs, & Marinazzo, 2016; Griffa, Baumann, Thiran, & Hagmann, 2013; Millan et al., 2012; Warren et al., 2014). Finally, replicating our results in different cohorts during the same period would be of paramount importance to ensure reproducibility.

## CONCLUSION

In this study, we used tools from network neuroscience and control theory to examine how the brain’s relatively rigid white matter architecture gives rise to a diverse repertoire of flexible neural dynamics, during normative development. We demonstrated that a brain region’s ability to display temporal flexibility in its functional expression positively correlated with the relative proportion of its anatomical edges reaching different cognitive modules across the brain. Indeed, this relationship was strongly mediated by the region’s boundary controllability, that is its capacity to integrate information across multiple cognitive processes. Overall, this work addresses the central question of how the human brain’s anatomical pathways support changes in flexible neural dynamics across late childhood, adolescence, and early adulthood, and provides a framework that could be used to study how neurological and psychiatric disorders emerge during that critical period of high neuroplasticity. Such analyses can leverage data on neurocognitive performance and clinical features available on this sample.

## AUTHOR CONTRIBUTIONS

Shi Gu: Conceptualization; Data curation; Analyses; Methodology; Visualization; Writing – original draft; Writing – review & editing. Panagiotis Fotiadis: Conceptualization; Data curation; Analyses; Methodology; Visualization; Writing – original draft; Writing – review & editing. Linden Parkes: Conceptualization; Visualization; Writing – review & editing. Cedric H. Xia: Conceptualization; Visualization; Writing – review & editing. David R. Roalf: Conceptualization; Writing - review & editing. Ruben C. Gur: Conceptualization; Writing - review & editing. Raquel E. Gur: Conceptualization; Writing - review & editing. Theodore D. Satterthwaite: Conceptualization; Supervision; Funding Acquisition; Writing - review & editing. Danielle S. Bassett: Conceptualization; Supervision; Funding Acquisition; Writing - review & editing.

## FUNDING INFORMATION AND CONFLICTS OF INTEREST

Linden Parkes acknowledges support from the Brain & Behavior Research Foundation (2020 NARSAD Young Investigator Grant). Ruben C. Gur and Raquel E. Gur acknowledge support from the National Institutes of Health (R01 MH119219) and the Lifespan Brain Institute of Penn Medicine and Children’s Hospital of Philadelphia. David R. Roalf acknowledges support from the National Institute of Mental Health (R01 MH119185, R01 MH120174) and the National Institute of Aging (R56 AG066656). Theodore D. Satterthwaite and Danielle S. Bassett together acknowledge support from the National Institutes of Health (R01 MH113550 and RF1 MH116920). Theodore D. Satterthwaite also acknowledges support through NIH R01 MH120482. The authors declare no conflicts of interest.

## CITATION DIVERSITY STATEMENT

Recent work in several fields of science has identified a bias in citation practices such that papers from women and other minority scholars are under-cited relative to the number of such papers in the field (Caplar, Tacchella, & Birrer, 2017; Dion, Sumner, & Mitchell, 2018; Dworkin et al., 2020; Maliniak, Powers, & Walter, 2013; Mitchell, Lange, & Brus, 2013). We obtained the predicted gender of the first and last author of each reference by using databases that store the probability of a first name being carried by a woman (Dworkin et al., 2020). By this measure (and excluding self-citations to the first and last authors of our current paper), our references contain 12.15% woman(first)/woman(last), 9.35% man/woman, 23.36% woman/man, and 55.14% man/man. This method is limited in that a) names, pronouns, and social media profiles used to construct the databases may not, in every case, be indicative of gender identity and b) it cannot account for intersex, non-binary, or transgender people. We look forward to future work that could help us better understand how to support equitable practices in science.

## Notes

### Competing Interest Statement

The authors have declared no competing interest.

